# Development and validation of a combined species SNP array for the European seabass (*Dicentrarchus labrax*) and gilthead seabream (*Sparus aurata*)

**DOI:** 10.1101/2020.12.17.423305

**Authors:** C. Peñaloza, T. Manousaki, R. Franch, A. Tsakogiannis, A. Sonesson, M. L. Aslam, F. Allal, L. Bargelloni, R. D. Houston, C. S. Tsigenopoulos

**Author notes:** Corresponding authors Ross Houston and Costas Tsigenopoulos. These authors contributed equally to this manuscript (C. Peñaloza, T. Manousaki, R. D. Houston, C. S. Tsigenopoulos).

## Abstract

SNP arrays are powerful tools for high-resolution studies of the genetic basis of complex traits, facilitating both population genomic and selective breeding research. The European seabass (*Dicentrarchus labrax*) and the gilthead seabream (*Sparus aurata*) are the two most important fish species for Mediterranean aquaculture. While selective breeding programmes increasingly underpin stocky supply for this industry, genomic selection is not yet widespread. Genomic selection has major potential to expedite genetic gain, in particular for traits practically impossible to measure on selection candidates, such as disease resistance and fillet yield. The aim of our study was to design a combined-species 60K SNP array for both European seabass and gilthead seabream, and to validate its performance on farmed and wild populations from numerous locations throughout the species range. To achieve this, high coverage Illumina whole genome sequencing of pooled samples was performed for 24 populations of European seabass and 27 populations of gilthead seabream. This resulted in a database of ~20 million SNPs per species, which were then filtered to identify high-quality variants and create the final set for the development of the ‘MedFish’ SNP array. The array was then tested by genotyping a subset of the discovery populations and demonstrated a high conversion rate to functioning polymorphic assays on the array (92% in seabass: 89% in seabream) and repeatability (99.4 - 99.7%). The platform interrogates ~30K markers in each fish species, includes features such as SNPs previously shown to be associated with performance traits, and is enriched for SNPs predicted to alter protein function. The array was demonstrated to be effective at detecting population structure across a wide range of fish populations from diverse geographical origins, and to examine the extent of haplotype sharing among Mediterranean fish farms. Therefore, the MedFish array enables efficient and accurate high-throughput genotyping for genome-wide distributed SNPs on each fish species, and will facilitate stock management, population genomics approaches, and acceleration of selective breeding through genomic selection.

## 1. Introduction

Modern aquaculture selective breeding is embracing modern sequencing and genotyping technologies via the inclusion of genomic information to sustainably increase genetic gain. The same genomic tools can facilitate improvements to methods for forming base populations for breeding programmes by computing well-characterized genetic variability and relationships, which is important for many aquaculture species still in the process of domestication [1, 2]. To achieve these goals in target species typically requires the generation of genome-wide genetic marker data (usually SNP markers) across large numbers of individuals. When paired with trait recording on the genotyped individuals, such datasets can be applied to examine the genetic architecture of production traits of interest, including detection of quantitative trait loci (QTL) using genome-wide association studies (GWAS). If the detected QTL are of sufficiently large effect, flanking markers can be utilized to select candidates with favourable alleles at the QTL, also known as Marker-Assisted Selection (MAS). While MAS has been successfully applied for a small number of traits, such as resistance to infectious pancreatic necrosis in Atlantic salmon [3, 4], most traits of interest for aquaculture are underpinned by a polygenic architecture [1, 5, 6]. For such traits, genome-wide SNP markers combined with phenotype data on a reference population can be used to estimate genomic breeding values for selection candidates [7]. Genomic selection is predicted to result in a notably higher selection accuracy and therefore genetic gain in aquaculture breeding programmes, as has also been demonstrated in early studies in several aquaculture species [1, 8], including European seabass (*Dicentrarchus labrax*) [9] and gilthead seabream (*Sparus aurata*) [10, 11].

The European seabass and the gilthead seabream are the two most important fish species in Mediterranean aquaculture. At the European level, they rank third and fourth, respectively, in terms of value after the Atlantic salmon and the rainbow trout [12]. Substantial genomic tools have been developed for both species, including the assembly and characterization of high-quality reference genomes [13, 14]. Medium or high density SNP arrays have been developed for several other important finfish aquaculture species such as rainbow trout (*Oncorhynchus mykiss*) [15], Atlantic salmon (*Salmo salar*) [16, 17], catfish (*Ictalurus furcatus and I. punctatus*) [18, 19], common carp (*Cyprinus carpio*) [20], Arctic charr (*Salvelinus alpinus*) [21], and Nile tilapia (*Oreochromis niloticus*) [22–24], which have been used for studies into population structure, genetic diversity, signatures of domestication, genetic architecture of traits of interest, and testing of genomic selection. A 57K SNP array was also recently developed for European sea bass [25] and has been applied to assess the genetic basis of resistance to viral nervous necrosis. However, while this array is available on request from the GeneSea consortium, there is a need for a publicly available SNP array for both European seabass and gilthead seabream.

Herein, we describe the generation of an extensive and comprehensive SNP database for European seabass and gilthead seabream across Europe by extensive sampling and pooled sequencing of ~25 populations per species from wild and aquaculture sites. From this SNP database, a subset of ~60K SNPs was chosen based on several filtering criteria to give thorough coverage of each species’ genome. The SNP array was created and validated on several of the discovery populations, including demonstration of its utility for detecting population structure and excess haplotype sharing between farmed populations. This open-access tool will provide new opportunities to the scientific community and industry for genome-scale research and application to improve selective breeding in these two focal European aquaculture species.

## 2. Materials and Methods

### 2.1. Samples for SNP discovery

A diverse range of farmed and wild populations of the European seabass (n = 24) and the gilthead seabream (n = 27) were collected for SNP discovery. A farmed population was defined as that composed of fish originating from the same commercial hatchery or established farm. A total of 538 European seabass individuals were sampled from 14 farmed and 10 wild populations distributed across the Mediterranean Sea, and a total of 642 gilthead seabream individuals were sampled from 12 farmed and 15 wild populations from the Mediterranean and the Atlantic (Table 1). Fin clips were collected from 15 to 30 individuals per population and stored in absolute ethanol until transportation to either The University of Edinburgh (UK), The Hellenic Centre for Marine Research (Greece) or The University of Padova (Italy) for DNA extraction.

**Table 1.** Summary of the European seabass and gilthead seabream populations sampled for sequencing and SNP discovery.

| Species | Origin | Region | Country | Pool ID | No individuals per pool |
| --- | --- | --- | --- | --- | --- |
| European seabass | farmed | Mediterranean | France | Sba_farm_1 | 12 |
|  |  |  | Spain | Sba_farm_2 | 25 |
|  |  |  | Spain | Sba_farm_3 | 25 |
|  |  |  | Italy | Sba_farm_4 | 25 |
|  |  |  | Croatia | Sba_farm_5 | 25 |
|  |  |  | Croatia | Sba_farm_6 | 25 |
|  |  |  | Greece | Sba_farm_7 | 25 |
|  |  |  | Greece | Sba_farm_8 | 25 |
|  |  |  | Greece | Sba_farm_9 | 25 |
|  |  |  | Greece | Sba_farm_10 | 25 |
|  |  |  | Greece | Sba_farm_11 | 25 |
|  |  |  | Greece | Sba_farm_12 | 23 |
|  |  |  | Cyprus | Sba_farm_13 | 25 |
|  |  |  | Egypt | Sba_farm_14 | 15 |
|  | wild | Mediterranean | France | Sba_wild_1 | 25 |
|  |  |  | Spain | Sba_wild_2 | 11 |
|  |  |  | Morocco | Sba_wild_3 | 25 |
|  |  |  | Italy | Sba_wild_4 | 25 |
|  |  |  | Croatia | Sba_wild_5 | 12 |
|  |  |  | Greece | Sba_wild_6 | 25 |
|  |  |  | Greece | Sba_wild_7 | 25 |
|  |  |  | Cyprus | Sba_wild_8 | 15 |
|  |  |  | Turkey | Sba_wild_9 | 25 |
|  |  |  | Turkey | Sba_wild_10 | 25 |
| gilthead seabream | farmed | Mediterranean | France | Sbr_farm_1 | 25 |
|  |  |  | Spain | Sbr_farm_2 | 25 |
|  |  |  | Spain | Sbr_farm_3 | 25 |
|  |  |  | Italy | Sbr_farm_4 | 25 |
|  |  |  | Croatia | Sbr_farm_5 | 25 |
|  |  |  | Greece | Sbr_farm_6 | 14 |
|  |  |  | Greece | Sbr_farm_7 | 13 |
|  |  |  | Greece | Sbr_farm_8 | 25 |
|  |  |  | Greece | Sbr_farm_9 | 25 |
|  |  |  | Greece | Sbr_farm_10 | 25 |
|  |  |  | Israel | Sbr_farm_11 | 25 |
|  |  |  | Egypt | Sbr_farm_12 | 15 |
|  | wild | Atlantic | France | Sbr_wild_1 | 25 |
|  |  |  | Spain | Sbr_wild_2 | 25 |
|  |  |  | Spain | Sbr_wild_3 | 25 |
|  |  | Mediterranean | Spain | Sbr_wild_4 | 25 |
|  |  |  | Spain | Sbr_wild_5 | 25 |
|  |  |  | Tunisia | Sbr_wild_6 | 25 |
|  |  |  | Italy | Sbr_wild_7 | 25 |
|  |  |  | Italy | Sbr_wild_8 | 25 |
|  |  |  | Greece | Sbr_wild_9 | 25 |
|  |  |  | Greece | Sbr_wild_10 | 25 |
|  |  |  | Greece | Sbr_wild_11 | 25 |
|  |  |  | Greece | Sbr_wild_12 | 25 |
|  |  |  | Greece | Sbr_wild_13 | 25 |
|  |  |  | Turkey | Sbr_wild_14 | 25 |
|  |  |  | Turkey | Sbr_wild_15 | 25 |

### 2.2. DNA extraction and pooling for sequencing

High quality genomic DNA was isolated from each fin-clip using a salt-based extraction method [26]. The integrity of the DNA extractions was assessed by performing an agarose gel electrophoresis. DNA purity was evaluated by using a NanoDrop ND-1000 (Thermo Fisher Scientific) spectrophotometer. The extracted DNA was quantified in duplicate using the fluorescent-based Qubit^®^ quantitation assay (Thermo Fisher Scientific, cat #Q32850). DNA stocks were diluted to 10-30 ng/ul and then combined at equimolar concentrations into pools of 11-25 individuals per population. The majority of populations had a sample size of 25, and for these populations DNA pools were prepared twice (technical replicates). For the remaining few populations with fewer individuals (6 and 3 populations in the European seabass and gilthead seabream, respectively), a single population pool was prepared.

### 2.3. Library construction and sequencing

Two sequencing facilities provided the library preparation and sequencing services – the Norwegian Sequencing Centre (NSC) (Oslo, Norway) and Edinburgh Genomics (University of Edinburgh, UK). Both facilities followed the TruSeq^®^ PCR-free library preparation protocol to generate sequencing libraries from the pooled genomic DNA samples. Almost all European seabass population pools were sequenced on a HiSeq 4000 instrument (2x 150 bp) at NSC, whereas all gilthead seabream pools were sequenced on a HiSeq X Ten platform (2x 150 bp) at Edinburgh Genomics.

### 2.4. Bioinformatics analysis for SNP discovery

The sequencing reads of the pooled DNA samples were processed separately for each species using identical software and parameter values. Raw sequencing data were filtered using the fastp software v 0.20.0 [27]. Reads with a minimum length of 80 bp for which less than 20% of their bases showed a BQ≤20 were retained. Cleaned paired-end reads were then aligned to either the European seabass [13] or the gilthead seabream [14] genome assemblies using BWA v 0.7.8 [28]. Only primary alignments to the relevant reference genome were retained for further analysis. PCR duplicates were removed from the alignment files using SAMtools v 1.6 [29]. Alignment files were sorted and indexed using BWA. Variants were called separately for each species across all population pools using Freebayes v 1.20 [30] with GNU Parallel [31]. Freebayes was set to call a variant if either (i) a minimum of 3 reads supporting the non-reference allele was observed, or (ii) the allele frequency in the pool was above 0.05, after excluding alignments with a MQ<20.

This initial list of variants was then filtered using vcflib (https://github.com/vcflib/vcflib) to keep bi-allelic SNPs that (i) showed supporting reads on both strands, (ii) a sequence coverage ranging from 17X to 90X for the European seabass and from 25X-100X for the gilthead seabream, (iii) at least two reads “balanced” to each side of the variant site, (iv) >90% of the observed alternate and reference alleles supported by properly paired reads, and (v) the ratio of mapping qualities between reference and alternate allele was between 0.9 and 1.1. SNPs were retained only if they had no interfering polymorphic sites within less than 35 bp upstream and downstream of the variant. The purpose of this filter was to identify markers compatible with array design and eliminate SNPs that could fail the assay due to flanking polymorphism interfering with probe annealing. The minor allele frequency (MAF) was estimated for all SNPs that were successfully genotyped in more than 18 population pools per species, after averaging the estimated MAF of the technical replicates. To avoid spurious SNPs resulting from sequence differences between paralogues, only SNPs with a MAF between 0.05 - 0.45 were retained for further SNP selection. From this list of candidate markers, 35 bp probes were extracted downstream and upstream from each SNP. The 71-mer nucleotide sequences were then submitted to Thermo Fisher for further quality check and *in silico* probe scoring.

### 2.5. SNP selection

As an initial filtering step, and as recommended by Thermo Fisher Scientific, the remaining SNPs were filtered to avoid *A/T* and *C/G* polymorphisms because they require twice the number of probes for genotyping compared to other types of SNP polymorphisms. The remaining SNPs were divided into selection tiers and were sequentially included in the MedFish platform based on the following hierarchy of importance.

First, SNPs were included as high priority markers based on evidence of their association with relevant production traits. For the European seabass, markers associated with mandibular prognathism [32], resistance to viral nervous necrosis [9], and sex [33] were included. For the gilthead seabream, the set of markers of this type comprised SNPs associated with production traits of high economic importance – i.e., fat content, weight, tag weight and length to width ratio [34] – and resistance to photobacteriosis [11]. Importantly, if the aforementioned SNPs were not identified through our pool-sequencing experiment, they were not included directly on the platform. Instead, the economically relevant marker was substituted by a proxy SNP that was chosen by screening the surrounding region for the closest high quality variant present in our dataset.

A second group of SNPs included in the MedFish SNP array is shared with other platforms that were developed in parallel at the time by the GeneSea consortium [25]. The purpose of including a subset of markers from the existing platforms was to facilitate backward compatibility and cross-study comparison, especially via the use of genotype imputation.

A third criterion for inclusion of SNPs on the MedFish platform was based on their predicted effect on protein coding genes. SNPs on genes may affect protein function, for example, by causing truncated proteins. To potentially target variants with a potential functional effect, which may have a direct impact on relevant phenotypes, the list of high confidence variants identified in the European seabass and the gilthead seabream genomes were annotated with SNPEff v 4.3 [35]. For both species, SNPs that were predicted to have a HIGH functional effect on proteins were considered important and included as high priority markers in the array.

Fourthly, from the total number of ~1.1 million SNPs per fish species that were submitted as 71-mers to Thermo Fisher for *in silico* probe evaluation, only those that were categorised as either ‘recommended’ or ‘neutral’ became the pool from which array SNPs were selected. From the substantial SNP database generated in this study, markers were selected to achieve good coverage of the reference genomes of the European seabass and gilthead seabream following [33]. In brief, markers were selected along each fish chromosome at a variable density depending on the estimated local nucleotide diversity (π), as in European seabass [13] and other fish species [36] a positive correlation between nucleotide diversity and recombination rate has been observed. For SNPs that were mapped to the “UN” chromosome of the European seabass, the synthetic chromosome was split into contigs that had been previously concatenated by 100 Ns. The contigs were isolated and the SNPs that were located within them where remapped with the contigs starting position set to 1. The genomes of both fish species were divided into 70 kb (for European seabass) or 85 kb (for gilthead seabream) non-overlapping windows and local nucleotide diversity was estimated with VCFtools v 0.1.15 [37]. Genomic windows were categorized into one of the following classes depending on their estimated π value: π ≤0.001 (Class 1), 0.001< π ≤0.002 (Class 2), 0.002< π ≤0.003 (Class 3), 0.003< π ≤0.004 (Class 4) and π >0.004 (Class 5). SNPs were chosen to cover all chromosomes of both fish species with a variable SNP density – ranging from 1-5 SNPs – depending on the diversity class of each region. For the SNP selection process carried out within each type of diversity class window, two factors were considered as the main inclusion criteria: (i) the MAF for the SNPs in the window and (ii) the physical distance between markers. All discovered markers were divided into three different MAF categories (>0.3, 0.3-0.2 and 0.2-0.1). SNP markers within the MAF >0.3 category were prioritized across all five window Classes such that at least 50% of the markers selected for each type of diversity window came from the most informative SNP category (Table 2). Within each window, SNPs were selected successively from each MAF category by requiring a minimum inter-marker distance of 10,000 bp with any other previously chosen set of markers. To fill the remaining target of ~30K SNPs per species, the physical distance between pairs of pre-selected SNPs was calculated, and intervals then sorted by length in decreasing order. SNP markers were then included sequentially (one SNP per interval) irrespective to its MAF.

**Table 2.** Summary of SNP selection approach. A variable number of SNPs was selected along chromosomes according to the local nucleotide diversity (π) estimates for non-overlapping genomic windows

| Genomic window Diversity Class | Range | No of SNPs sampled per window | No SNPs sampled per allele frequency category per window |  |  |
| --- | --- | --- | --- | --- | --- |
| | | | $>0.3$ | $0.2-0.3$ | $0.1-0.2$ |
| Class 1 | $\pi \leq 0.001$ | 1 | 1 | 0 | 0 |
| Class 2 | $0.001 < \pi \leq 0.002$ | 2 | 2 | 0 | 0 |
| Class 3 | $0.002 < \pi \leq 0.003$ | 3 | 2 | 1 | 0 |
| Class 4 | $0.003 < \pi \leq 0.004$ | 4 | 2 | 2 | 0 |
| Class 5 | $\pi > 0.004$ | 5 | 2 | 2 | 1 |

A final list of ~70K SNPs was sent to Thermo Fisher for the creation of the 60K SNP array. This 384-format genotyping array was called the MedFish array, reflecting the two European Union funded consortium projects MedAID and PerformFish (see the Acknowledgements section for details).

## 3. Validation of the MedFish array through population genomic analyses

### 3.1. Genotyping

A subset of 502 European seabass and 478 gilthead seabream fin clips from the same populations used for SNP discovery were sent to IdentiGEN (Ireland) for DNA extraction and genotyping with the MedFish SNP array (Table 3). To assess the repeatability and quantify the putative error rate of the platform, a single replicate sample (one per species) was genotyped twelve times across three different arrays. The proportion of SNPs at which the (replicate) individuals shared identical-by-state alleles was calculated.

**Table 3.** Fish samples genotyped using the combined species MedFish SNP array.

| Species | Origin | Population ID | Country | No fish typed | No fish passing QC |
| --- | --- | --- | --- | --- | --- |
| European seabass | farmed | Sba_farm_1 | France | 12 | 12 |
|  |  | Sba_farm_2 | Spain | 25 | 25 |
|  |  | Sba_farm_3 | Spain | 25 | 21 |
|  |  | Sba_farm_4 | Italy | 24 | 24 |
|  |  | Sba_farm_6 | Croatia | 25 | 24 |
|  |  | Sba_farm_7 | Greece | 25 | 21 |
|  |  | Sba_farm_8 | Greece | 25 | 21 |
|  |  | Sba_farm_9 | Greece | 24 | 21 |
|  |  | Sba_farm_10 | Greece | 25 | 23 |
|  |  | Sba_farm_11 | Greece | 24 | 21 |
|  |  | Sba_farm_12 | Greece | 23 | 22 |
|  |  | Sba_farm_13 | Cyprus | 23 | 21 |
|  |  | Sba_farm_14 | Egypt | 14 | 12 |
|  | wild | Sba_wild_Mediterranean |  | 208 | 192 |
| gilthead seabream | farmed | Sbr_farm_1 | France | 24 | 20 |
|  |  | Sbr_farm_2 | Spain | 18 | 17 |
|  |  | Sbr_farm_3 | Spain | 25 | 25 |
|  |  | Sbr_farm_5 | Croatia | 19 | 19 |
|  |  | Sbr_farm_6 | Greece | 13 | 12 |
|  |  | Sbr_farm_7 | Greece | 13 | 13 |
|  |  | Sbr_farm_8 | Greece | 21 | 21 |
|  |  | Sbr_farm_9 | Greece | 24 | 24 |
|  |  | Sbr_farm_10 | Greece | 20 | 19 |
|  |  | Sbr_farm_11 | Israel | 13 | 13 |
|  |  | Sbr_farm_12 | Egypt | 15 | 14 |
|  | wild | Sbr_wild_Atlantic |  | 28 | 28 |
|  |  | Sbr_wild_Mediterranean |  | 245 | 227 |

SNP quality control and genotype calling from the intensity files was performed using the Axiom Analysis Suite software v 2.0.035 at default parameter values for diploid species (call rate (CR) > 97; dish QC (DQC) >0.82). Because a significant fraction of the European seabass samples had a CR below the default value of 97 (201 individuals), the threshold was reduced to 93, allowing to recover genotypes for 460 individual samples.

### 3.2. Population structure

The combined species ~60K MedFish array was tested by performing a principal component analysis (PCA) on the genotypes of a wide range of Mediterranean (and a few Atlantic) European seabass and gilthead seabream farmed and wild population typed with the platform. These individuals were part of the same set of samples used for the SNP discovery process from Pool-seq data (Table 3). A QC-filtered SNP dataset was created by applying the following filters in PLINK v2.0 [38]. Bi-allelic SNPs were retained for analysis if they had (i) a call rate > 0.95, (ii) MAF > 0.01, (iii) HWE test p-value ≥ 1e-4 (estimated separately for each population) and (iv) no pairs with a squared LD correlation (r^2^) > 0.2 occurred within a 100 kb window. For duplicated or related individuals with a kinship coefficient (KING-rob) > 0.177 (first-degree relatives or closer) only one member of a pair was retained for further analysis. All individuals to be evaluated required having < 10% missing genotypes. The structure of the genotyped populations was investigated using the R package LEA [39].

### 3.3. Analysis of haplotype sharing

To assess the ability of the SNP array to identify historical connections between farmed populations, a haplotype sharing analysis was performed on the farmed population samples (13 European seabass farms; 11 gilthead seabream farms). A SNP dataset in which all individual and SNP QC filters had been applied (see Population structure section), except the removal of markers based on linkage disequilibrium (measured as r^2^) was used for the analysis. Markers that were not located on chromosomes of the reference genome assemblies were removed from the dataset. Haplotypes were inferred for each individual using the software fastPHASE v1.4.8 [40]. All individuals were phased together in a single analysis, taking into consideration their population labels during the model fitting procedure. For both fish species, the number of random starts of the EM algorithm (T) was set to 20, the number of iterations (C) was set to 35, and the number of haplotype clusters (K) to 8.

The reference genomes of both species were divided in 1 Mb non-overlapping windows using BEDTools v2.25 [41]. SNP-based haplotype variants were defined for each window. The last window of each chromosome was excluded from the analysis. Since the number of haplotypes can be influenced by sample size, the same number of individuals were randomly chosen from each farmed population (6 individuals for the European seabass and 9 for the gilthead seabream). For each individual within a farm, the two haplotypes at any given locus were used to screen the whole dataset for an exact match. All matches with other individuals from a different farm were recorded. The totals were then summed across all individuals that belonged to the same farm, and the proportion of shared haplotypes across farms calculated.

To assess whether a pair of farms had excess haplotype sharing, 1,000 permutations were performed. For each permutation, all individuals from a fish species were randomly assigned to an arbitrary farm.

### 3.4. Ethics statement

The fish fin clip collected in this study were obtained from commercial samples or specific sampling efforts managed and sampled always in accordance with the European directive 2010/63/UE on the protection of animals used for scientific purposes.

## 4. Results

### 4.1. SNP array development

The pooled DNA sequencing of 24 European seabass and 27 gilthead seabream populations produced 8,205 and 23,784 million paired-end reads, respectively. The alignment of the post-quality filtered reads against each species reference genome resulted in the discovery of ~17 million polymorphisms in the European seabass and ~34 million putative polymorphisms in the gilthead seabream genomes (including both SNPs and indels). The generated sequence led to an average coverage at SNP variant sites of 36X in the European seabass and 63X in the gilthead seabream. After applying the QC filters on the variant call set (see Materials and Methods), a pool of 1,056,218 and 1,015,264 high confidence SNPs in the European seabass and the gilthead seabream, respectively, remained for SNP selection. The QC filter that removed the largest amount of data was the restriction to keeping variants without polymorphisms in close proximity (within 35 bp on either side). This filter alone removed 88% and 96% of the variants discovered *de novo* in the European seabass and gilthead seabream, respectively.

Following the submission and evaluation of these filtered SNPs by Thermo Fisher, SNPs were sampled along chromosomes following the strategy of selecting a number of SNPs in proportion to the putative local recombination rate (measured as π) of the genomic region. Notably, and in comparison to the European seabass, a particularly high number of polymorphisms were initially discovered along the chromosomes of the gilthead seabream, particularly towards the terminal ends of the chromosome-level scaffolds. The most likely cause was that in this species the higher average sequencing depth of 63X (compared to 36X in the European seabass population pools) enabled the discovery of variants segregating at a lower frequency. Consequently, when the QC filter that removed SNPs with interfering markers in close proximity was applied to the gilthead seabream dataset, a substantial number of markers were filtered-out from regions of the genome exhibiting high levels of genetic polymorphism. This led to fewer SNPs left to choose from for assay design in regions of the gilthead seabream genome that showed putative higher recombination (e.g. chromosome ends) and for which a higher number of SNPs had to be sampled, based on our SNP selection strategy. Therefore, the SNP selection strategy led to a more even sampling of SNPs along the gilthead seabream genome. While in European seabass, the array SNPs followed the expected pattern of the SNP selection strategy, with more markers being assayed towards the terminal ends of the chromosome-level scaffolds (Figure 1).

**Figure 1.**
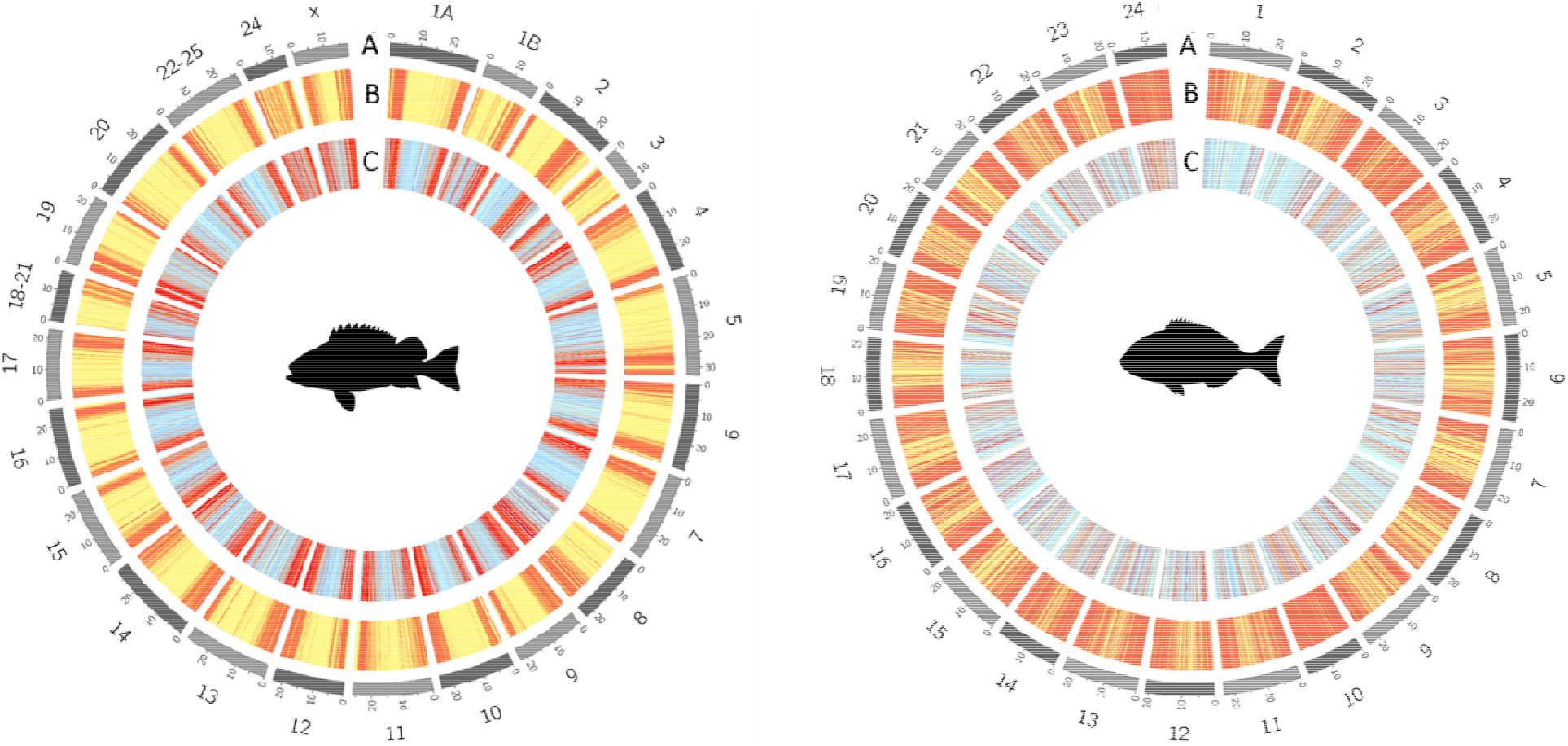
Schematic representation of the distribution of array markers in the European seabass (left) and gilthead seabream (right) genomes after following a SNP selection strategy based on local nucleotide diversity. (A) Chromosome number. (B) Levels of diversity (π) estimated over 70 kb and 85 kb windows in the European seabass and gilthead seabream, respectively. Red bars represent regions with high nucleotide diversity. (C) Genome-wide distribution of markers on the combined-species SNP chip. Light blue bars represent windows for which 1-3 SNPs were selected. Red bars represent windows for which more than four SNPs were selected.

The final MedFish SNP array was designed to interrogate 29,888 SNPs in the European seabass genome and 29,807 SNPs in the gilthead seabream genome. Among these markers, 4,560 SNPs (15%) in the European seabass and 3,208 SNPs (11%) in the gilthead seabream are shared with other platforms that were being developed at the time of this study [25]. A significant fraction of the SNPs on the platform are located in genes (46% seabass; 32% seabream), among which 107 and 179 SNPs, respectively, were predicted *in silico* to have high functional effects on proteins. For the SNPs included on the array, the physical distance between consecutive markers was similar for both species and averaged 20 kb in the European seabass and 19 kb in the gilthead seabream. The largest gaps between markers (200-300 kb) represented a small fraction of the platform and comprised five regions on chromosomes 1, 4, 9, 13 and 16 of the gilthead seabream. Detailed examination revealed that these regions lacked suitable markers matching our SNP selection criteria. No large regions in the European seabass genome were devoid of assays, with the highest inter-marker distance being ~120 kb.

Two metrics were used to assess the performance of the assays on the array: (i) conversion rate and (ii) platform error rate. Here the conversion rate is defined to be the fraction of probes that yielded strong signals with high-quality clusters discerning different genotypes. The conversion rate of the European seabass fraction of the array was 91.9%, whereas the gilthead seabream assays on the array had a conversion rate of 88.7% (Table 4). In terms of the informativeness of the markers on the platform, for 99.8% of the validated loci in the European seabass the MAF was >5%. In the case of the gilthead seabream, 98.7% of the markers had a MAF>5%. The process of calculating the platform error rate involved genotyping two samples (one per species) twelve times each. For European seabass, one replicate sample failed to generate a CEL file, consequently eleven samples remained for evaluation. The repeatability of the assays was 99.43% for the European seabass and 99.75% for the gilthead seabream. Taken together these metrics support the high quality and reliability of the genotype data generated by the MedFish SNP array.

**Table 4.** Number of SNPs for each species falling within each Axiom quality class. The categories are based on cluster properties and QC metrics.

| **Conversion type\*** | **No European seabass (%)** | **No gilthead seabream (%)** |
| --- | --- | --- |
| Polymorphic high resolution | 26,466 (88.55%) | 26,369 (88.47%) |
| No minor homozygote | 993 (3.32%) | 75 (0.25%) |
| **Total high quality polymorphic** | **27,459 (91.87%)** | **26,444 (88.72%)** |
| Monomorphic high resolution | 26 (0.09%) | 36 (0.12%) |
| Off-target-variant (OTV) | 50 (0.17%) | 78 (0.26%) |
| Call rate below threshold (97%) | 889 (2.97%) | 1,292 (4.33%) |
| Other | 1,464 (4.90%) | 1,957 (6.57%) |
| **Total SNPs on the array** | **29,888 (100%)** | **29,807 (100%)** |

\* The Conversion type follows Thermo Fisher's terminology:
*PolyHighResolution* = Class with the highest quality probes. SNP is polymorphic and the presence of both the major and minor homozygous clusters is observed.
*NoMinorHom* = similar to a *PolyHighResolution*, but no evidence of individuals with minor homozygous genotypes, presumably due to a low genotype frequency.
*MonoHighResolution* = SNP can reliably be scored as monomorphic.
*Off-target variant (OTV)* = SNPs where additional (i.e. more than three) clusters are observed, making genotype calling ambiguous.
*CallRateBelowThreshold* = SNP with the expected number of clusters (usually 3, one for each possible genotype), but where the proportion of samples scored at the SNP falls below a user-defined threshold.
*Other* = SNPs that do not fall in any of the above categories.

### 4.2. Population structure

To gain a general overview on the population structure within each species we performed a PCA analysis on the genotyping data. The two first principal components (PCs) explained 21% and 11% of the total variance for European seabass and gilthead seabream, respectively (Figure 2).

**Figure 2.**
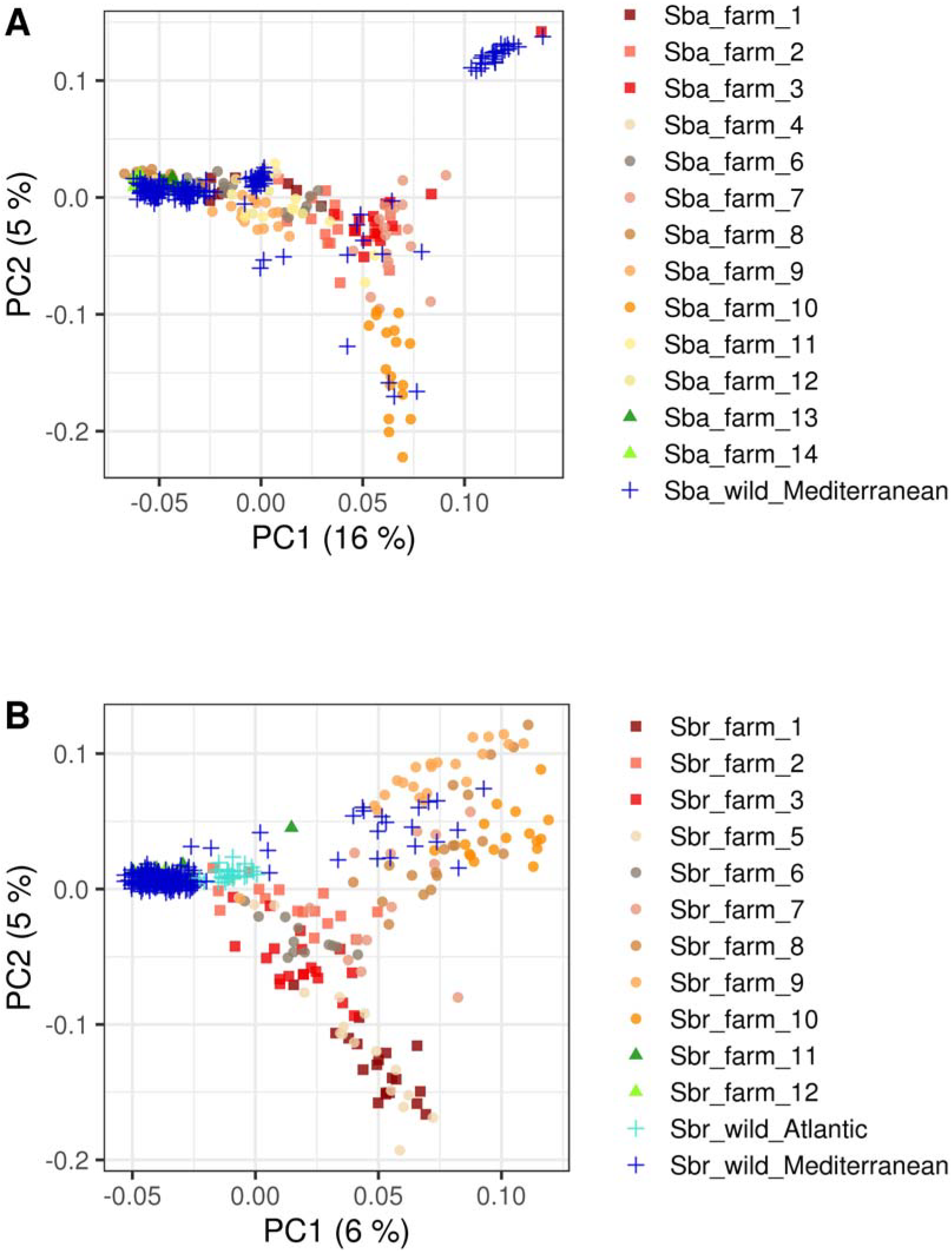
Population genetic structure of farmed and wild European seabass and gilthead seabream populations. (A) PCA of 460 European seabass individuals from fourteen Mediterranean populations. All wild populations are from the Mediterranean and are grouped under the same population label. (B) PCA of 478 gilthead seabream individuals from thirteen populations. Wild individuals are grouped by origin into either a Mediterranean or Atlantic population. The different point symbols separate samples by origin in (i) farms from the West of the Mediterranean (▪), (ii) farms from the Centre of the Mediterranean (⋅), and (iii) farms from the East of the Mediterranean (▴), from (iv) wild populations (✝).

In European seabass, most of the sampled farmed populations form a loose cluster along PC1, which explains 16% of the variance. No geographical cline is observed as farms from the West (France, Spain), Centre (Italy, Croatia, Greece) and East (Cyprus, Egypt) of the Mediterranean cluster at least partially in this dimension. On the other hand, three distinctive clusters are recognized for the wild European seabass populations, with a few exceptions corresponding to individuals clustering near farmed populations instead. PC2 explains 5% of the total variation and mainly separates (i) a single well-defined wild population cluster, (ii) a large group containing most of the farmed and wild seabass populations, and (iii) a group of individuals that belong to a farm sampled from the Centre of the Mediterranean (farm № 10 sampled from a Greek hatchery) (Figure 2A).

Regarding the gilthead seabream, the sampled farmed populations form a continuum along PC1 rather than discrete units. Although the majority of gilthead seabream wild populations were sampled from the Mediterranean Sea, a few populations from the Atlantic coast of France and Spain were included in the analysis. Individuals sampled from a wide range of wild populations group by origin into either a Mediterranean or Atlantic cluster on one extreme of the PC1 axis. While individuals sampled from farms from the Centre of the Mediterranean (either Italy or Greece) are represented at the other end of the PC1 axis. PC2 accounts for 5% of the variance and distinguishes two groups of overlapping farmed populations that partially coincide with their macro-region of origin. The first group is composed only of farms located in the Centre of the Mediterranean (i.e. from either Italy or Greece). A few wild gilthead seabream individuals co-localize with this group of farmed samples. The second group is comprised of a mixture of all three farms sampled from the West of the Mediterranean (i.e. from either France or Spain) and a few populations sampled from farms located in the Centre of the Mediterranean, namely farm № 5 (from Croatia) and № 6 (from Greece) (Figure 2B).

### 4.3. Haplotype sharing analysis

After applying QC filters, a total of 21,822 SNPs in European seabass and 24,765 SNPs in the gilthead seabream remained for the assessment of haplotype sharing between pairs of Mediterranean fish farms.

The pairwise comparison among European seabass farms revealed that all populations showed an excess of haplotype sharing with at least one other Mediterranean farm (Fig. 3 A). A pairwise comparison of two Greek seabass farms (farm № 8 vs. farm № 12) resulted in the highest percentage of haplotype sharing (43%). The reverse relationship between these two farms (i.e. farm № 12 vs. farm № 8) is also significant but is ranked 9^th^ (19%) in terms of haplotype-sharing percentage among farms. This difference in reciprocal comparisons is explained by differences in the total numbers of shared haplotypes identified within each farm (File S1). Haplotypes from individuals of a European seabass farm located in Greece (farm № 7) were present at significant frequencies in all farms sampled from the West of the Mediterranean (i.e. farms of French or Spanish origin) and most of the seabass farms sampled from the Centre of the Mediterranean (i.e. either from Italy, Croatia or Greece). With regards to haplotype count (i.e. absolute number of haplotypes shared between farms), farm № 7 shares a significant number of haplotype variants with farms № 10 (hap = 1,466), № 2 (hap = 978) and № 3 (hap = 847).

**Figure 3.**
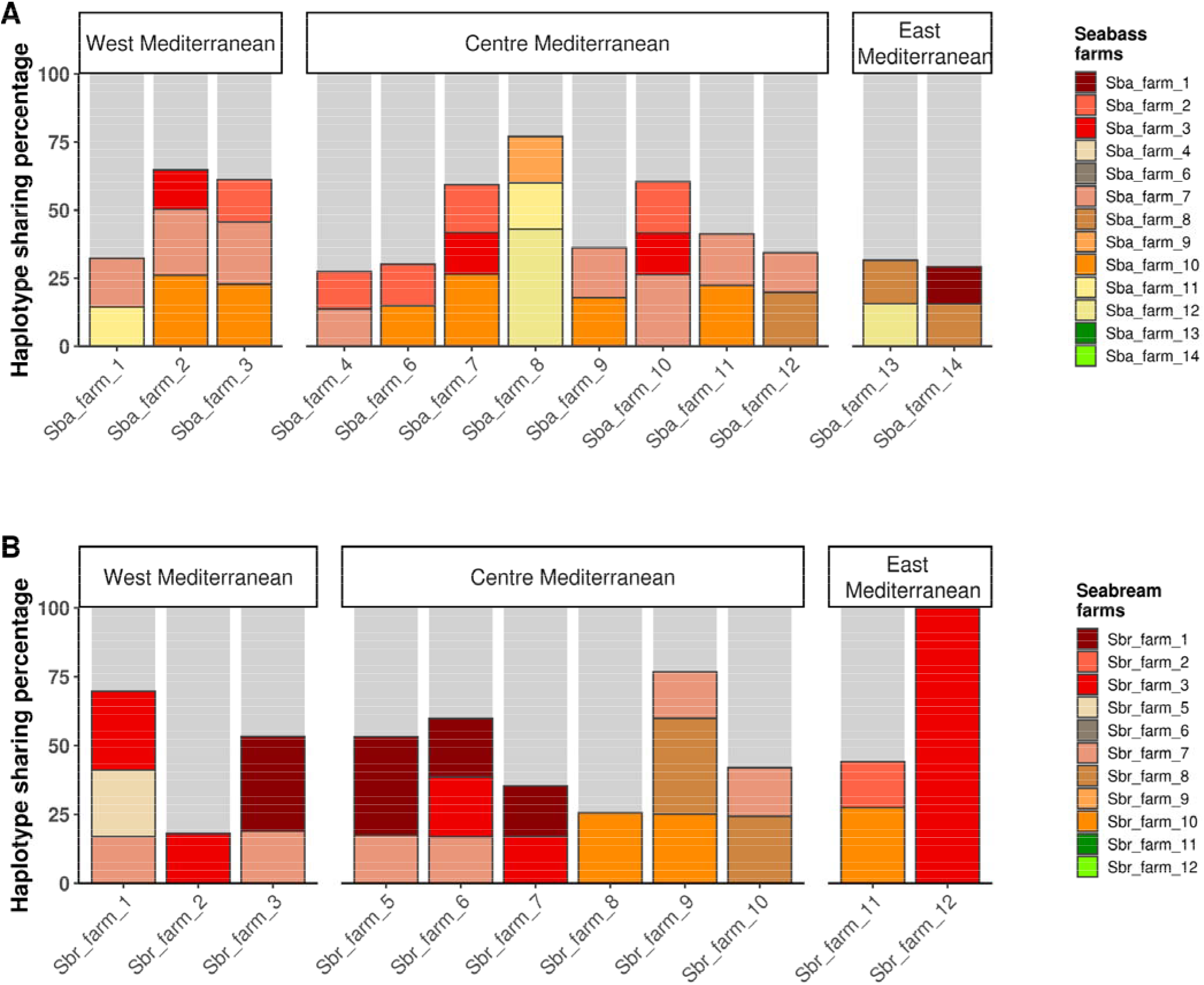
Haplotype sharing between farmed populations of (A) European seabass and (B) gilthead seabream. On the x-axes, the different farms sampled per species are stratified by geographical origin (either West, Centre or East Mediterranean). The stacked bar charts show the percentage of haplotypes (relative to the total number of shared haplotype variants identified in each farm) shared with another farm (y-axes), color-coded following the legend on the right. Only pairs of farm showing a statistically significant excess of shared haplotypes (p-value < 0.05) are shown.

In common with the results observed for European seabass, all gilthead seabream farms evaluated show an excessive sharing of haplotypes with at least one other Mediterranean farm (Fig. 3 B). In gilthead seabream, a clear break separates the farms from the West and Centre of the Mediterranean in two groups. One group includes six farms – farms № 1, 2, 3, 5, 6 and 7 – from diverse geographical origins (i.e. France, Spain, Croatia and Greece). The second group comprises farms exclusively based in Greece – farms № 8, 9 and 10. Reduced haplotype sharing was observed between the farmed populations from both aforementioned groups. Moreover, only one gilthead seabream farm – farm № 7 – had haplotypes that were also present at significant levels in farms belonging to both groups. A seabream farm sampled from the East of the Mediterranean (farm № 12) had the lowest total number of shared haplotypes among all commercial farms evaluated from both fish species. Most haplotypes identified in farm № 12 were unique and specific to the farm, which showed complete absence of shared variants with all but one Mediterranean farms (i.e. farm № 3).

## 5. Discussion

### 5.1. Properties of the combined species MedFish SNP array

A publicly available, combined species SNP chip that assays ~30K SNPs throughout the genome of two prominent Mediterranean fish species - the European seabass and the gilthead seabream – was developed. To evaluate the performance of the MedFish SNP array two metrics were analyzed: conversion rate and platform error rate. The conversion rate is a measure of the number of SNPs that are successfully assayed by a technology and reflects the quality of both the chosen SNPs and the technology used to score them [42]. Conversion rates were high for the SNP array regardless of the fish species. The assay conversion rate of the European seabass part of the array was 91.9%, while for the gilthead seabream part it was 88.8%. These values are slightly lower than terrestrial livestock species arrays (e.g. 92.6% in cattle and 97.4% in pigs), however, generally higher than those developed for aquatic organisms (e.g. 72.5% in oysters and 86.1% in catfish) [18, 43-45], comparable to the top performing finfish arrays [15, 25]. As a second metric to assess performance, the platform error rate was calculated based on the genotype concordance of repeated assays on the same individual. By this metric, the MedFish platform shows high genotype accuracy, with a repeatability ranging from 99.4% - 99.7%. This accuracy levels are comparable to those achieved with Illumina GoldenGate assays in humans (99.6%) or Affymetrix SNP chips in trout (99.4%) and pig (100%) [15, 43, 46]. Compared to other SNP arrays developed for aquaculture species, the MedFish platform stands out both in terms of genotype accuracy and repeatability. Until recently, high-throughput genotyping analysis was only achievable in these fish species by means of reduced-representation sequencing approaches [9, 11, 34]. Although a cost-efficient option, these techniques may suffer from inconsistent marker recovery across experiments and comparatively lower robustness to low quality input DNA [47]. Hence, the development of this combined species SNP array represents a powerful alternative for high-throughput genotyping in European seabass and gilthead seabream, facilitating molecular breeding applications, genetic stock identification and population and evolutionary studies in these emblematic fish species. Moreover, the fact that the two species are represented on the same platform increases the overall volume of arrays that can be purchased, which should reduce the cost of the array due to economy of scale. This reduced cost will be key to the uptake of the platform by aquaculture breeding and production companies for the routine application of genomic selection in a cost-effective way.

As part of the SNP chip design, over 25 farmed and wild populations (>500 individuals) per species were screened for highly informative markers. By following a DNA pooling approach, reliable genome-wide allele frequency information was obtained for several fish populations at a fraction of the effort of individual sequencing. Given the majority of the samples genotyped with the SNP array were also part of the SNP discovery process, metrics such as number and mean MAF of polymorphic markers reflected the performance of the SNP selection strategy. Despite the relatively small DNA pools (12-25 individuals), we were able to reliably identify and select informative markers for inclusion in our SNP array. The number of informative markers (MAF>0) was high for both fish species. For the European seabass, 23,900 SNPs (99.8%) of the validated markers were polymorphic, whereas for the gilthead seabream this type of markers comprised 26,017 SNPs (99.3%) of the data. The number of polymorphic markers was remarkably similar in wild and farmed populations of both species (90-99% across populations), demonstrating the efficacy of the SNP selection strategy for recovering highly informative markers in Pool-seq data sequenced at a high to moderate coverage across a wide range of different populations. When evaluating the MAF across European seabass and gilthead seabream populations, the allele frequency profiles were similar within species, and did not vary significantly by origin (either wild or farmed) (Fig S1). The mean MAF across the European seabass (0.33) and gilthead seabream (0.31) populations was higher than that reported when validating SNP arrays in Nile tilapia (0.29) and rainbow trout (0.25) [15, 22]. However, the high average MAF observed in this study is most likely influenced by the fact that most of the discovery populations were also used for the validation of the SNP chip. Nonetheless, it should be noted that the discovery population samples cover a large portion of the distribution range in the wild, and include the majority of commercial hatcheries for the two species.

A significant obstacle to the uptake of high-throughput genotyping technologies by the industry is the risk that a low fraction of a pre-built platform yields useful information. Indeed ascertainment bias is a common issue for genotyping arrays, and can be caused when designing platforms based on a reduced number of individuals [48]. Due to the fact that the MedFish 60K array was developed based on the screening of genetic data derived from an extensive sampling of dozens of Mediterranean fish populations and hundreds of fish of each species, it is tailored to maximize the retrieval of genetic information and provide an increased resolution for the analysis of farmed or wild stocks from this region.

### 5.2. Population structure and haplotype sharing analysis

To validate the MedFish SNP array, the genotyping data obtained from typing a diverse range of wild and farmed European seabass and gilthead seabream fish were used to perform a principal components (PC) and haplotype sharing analysis.

Regarding the European seabass populations, the two first PCs explained 21% of the genotypic variation. Interestingly, the wild Mediterranean populations span a continuum across the range of PC1, but has a rather smaller dispersal across the PC2 range. However, this continuum in PC1 has gaps and the wild populations seem to be divided into two clearly differentiated clusters, which may represent the two different lineages of European seabass described by Tine et al. [13]. All farmed European seabass populations have a more limited distribution across the range of the two PCs compared to the wild populations, forming clear clouds although not so dense as the wild populations, which is probably due to their smaller sample sizes (Table 3). Most farm populations fall within the range of the wild populations, with overlap among each other. Only a single farmed fish stock seems to have a more distinct pattern (Fig. 2 A; farm № 10), which might reflect either founder effects, stronger artificial selection, higher number of generations of selection, or any combination thereof. European seabass farms of different geographical origin tend to cluster together in the PC plot. For instance, farms № 2 and 3 (from Spain) group with farm № 7 (from Greece). This observation is consistent with the haplotype sharing analysis, as a significant number of 1 Mb SNP-based haplotype variants were jointly present in these farms. A high frequency of shared haplotypes between pairs of populations provides information about their historical relationship, reflecting either a common ancestry and/or gene flow between populations. In the context of aquaculture farming, a high frequency of shared haplotypes between farms might indicate (i) animal transfer between farms or (ii) the recent establishment of these farmed populations from the same wild source (i.e. recent population divergence). Since the PCA revealed that wild populations of European seabass form tight and distinctive clusters, it is likely that pairs of European seabass farms sharing a high frequency of haplotype variants are derived from human-mediated translocations of fishes. Another interesting finding is that few of the wild individuals fall clearly within the range of farm populations. This could be either due to greater genetic similarity of these farmed populations to certain wild populations that are poorly represented in the genotyped samples, or that these wild individuals are escapees from fish farms, a well-known phenomenon occurring in the Mediterranean [49, 50].

For the gilthead seabream populations, the PCA explained much less of the observed genetic variance (only 11% covered by both PC1 and PC2 summed up), showing a less clear structure for most of the populations sampled in this study. In this case, wild populations were sampled from two regions, the Mediterranean and Atlantic. Wild individuals segregate into two closely bound Mediterranean and Atlantic clusters, which is consistent with previous findings indicating a low genetic differentiation between basins [51]. Similar to European seabass, a few wild individuals are found scattered throughout farmed populations, likely representing escapees from local fish farms. Farmed gilthead seabream populations seem to be much more differentiated compared to their wild counterparts, with two broader clusters forming a gradient of overlapping farmed populations. The first group is composed only of farms located in Greece. The second cluster groups a mixture of all three farms sampled from the West of the Mediterranean (either France or Spain) and a few Greek farms (Figure 2B). This pattern may reflect artificial selection and/or different degrees of admixture between farms. The haplotype sharing analysis mirrors this finding and reinforces the idea that most seabream farms from the Mediterranean separate in two clusters, between which a reduced recent contact is observed. However, while the results for both the European seabass and gilthead seabream highlight the utility of the SNP array for detecting and studying population structure, more extensive studies are required to further assess these phenomena in the two species.

## 6. Conclusions

A medium density SNP array suitable for genotyping both the European seabass and the gilthead seabream was developed. The MedFish SNP array has a high proportion of functional and validated SNP assays, as demonstrated by its conversion rate (92% in the European seabass: 89% in the gilthead seabream) and repeatability (99.4 - 99.7%). The platform interrogates ~30K markers in each fish species, and includes features such as SNPs previously shown to be associated with performance traits and enrichment for SNPs predicted to have high functional effects on proteins. The SNP array was highly informative when tested on the majority of the discovery population samples, and was further validated by performing a population structure and haplotype sharing analysis across a wide range of fish populations from diverse geographical backgrounds. This recently developed platform will allow the efficient and accurate high-throughput genotyping of ~30K SNPs across the genomes of each fish species, facilitating population genomic research and the application of genomic selection for acceleration of genetic improvement in European seabass and gilthead seabream breeding programs.

## Data availability

Raw sequence reads from the European seabass and gilthead seabream population pools analyzed for SNP discovery have been deposited in NCBI’s Sequence Read Archive (SRA) under accession number PRJEB40423. Details of the allele frequencies of the SNPs on the MedFish array can be found in the Mendeley Data Repository (http://dx.doi.org/10.17632/k3rr2k7svk.1).

## Acknowledgements

This study was made possible by the EU projects MedAID (H2020 grant agreement No 727315) and PerformFish (H2020 grant agreement No 727610). We are grateful to both projects’ partners that contributed with samples for SNP discovery as well as the projects’ coordinators (A. López-Francos, B. Basurco, D. Furones and K. Moutou) and the H2020 project officer for enhancing a fruitful interaction between research consortia. The authors would also like to thank Christos Palaiokostas and Sara Faggion for providing additional information for trait-associated variants included in the SNP array. CP and RH are supported by funding from the BBSRC Institute Strategic Programme Grants BB/P013759/1 and BB/P013740/1. We also give thanks to the sequencing facilities Edinburgh Genomics (University of Edinburgh, UK) and the Norwegian Sequencing Centre (University of Oslo, Norway). TM would like to thank George Nomikos for valuable support on data analysis. This research was supported through computational resources provided by IMBBC (Institute of Marine Biology, Biotechnology, and Aquaculture) of the HCMR (Hellenic Centre for Marine Research). Funding for establishing the IMBBC HPC has been received by the MARBIGEN (EU Regpot) project, LifeWatchGreece RI, and the CMBR (Centre for the study and sustainable exploitation of Marine Biological Resources) RI. We also thank the GeneSea consortium (n° R FEA 4700 16 FA 100 0005) for providing some variants allowing interoperability with existing arrays.

