## Supplementary Figure 1 for "Development and validation of a combined species SNP array for the European seabass (*Dicentrarchus labrax*) and gilthead seabream (*Sparus aurata*)"

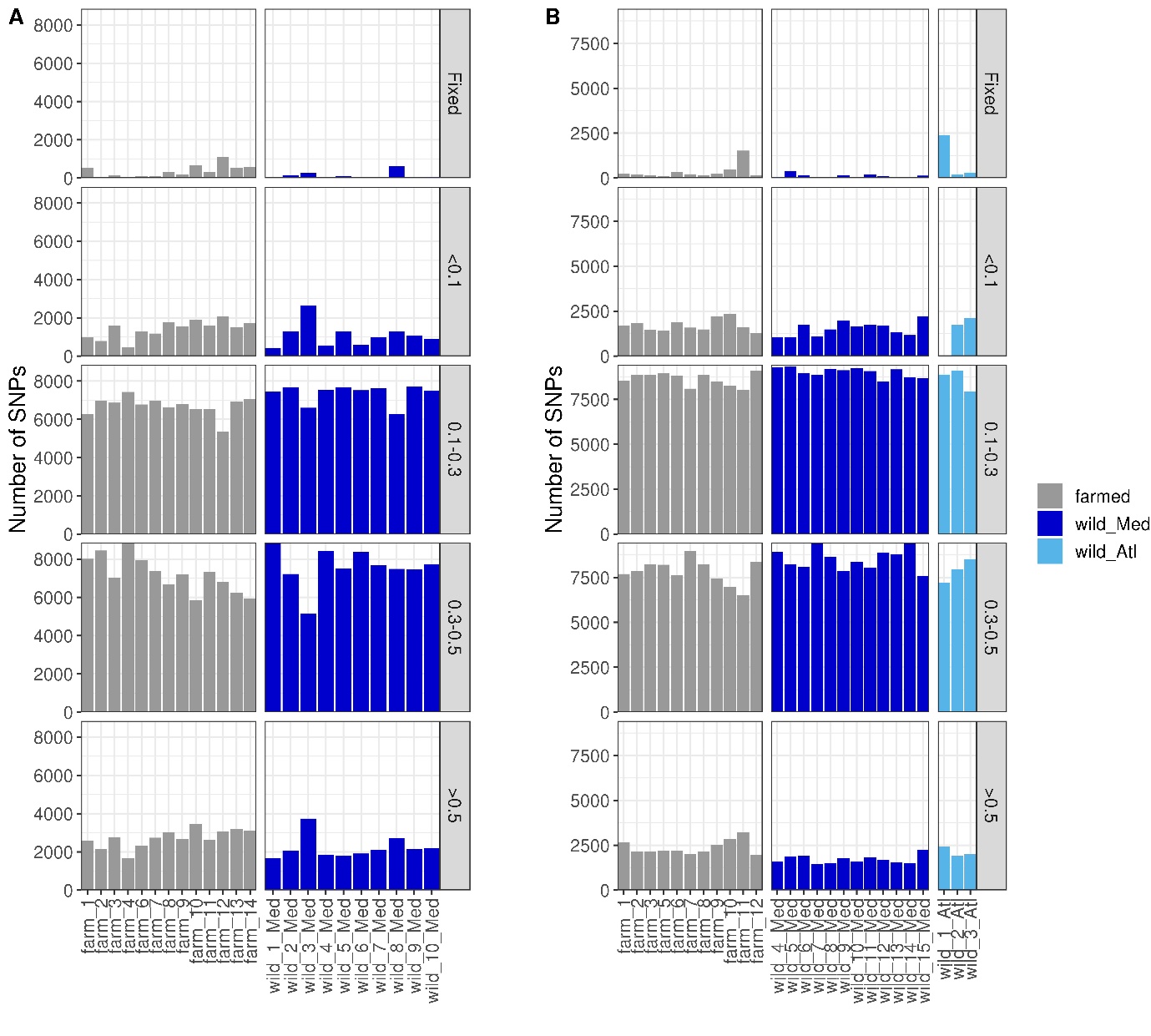

**Figure S1. Allele frequency of the SNPs typed with the combined species SNP array across different farmed and wild (A) European sea bass and (B) gilthead sea bream populations.** Panels are stratified by allele frequency as indicated in the labels along the y-axes on the right side. Populations are coloured coded according to their known origin as shown in the legend at the right.
